# Identifying Effective Cryoprotection Agents for Non-Model Bacterial Species

**DOI:** 10.64898/2025.12.04.692406

**Authors:** Ruth Wright, Bhuwan Abbot, Taylor Yonemura, Morgan Carter

## Abstract

Host-associated bacteria live amongst eukaryotes within varied niches and form relationships ranging from facultative to obligate. With advancement in studies of such symbiotic associations, fastidious bacteria are increasingly becoming targets for genetic manipulation. However, there are limited resources for screening possible agents enabling *in vitro* culturing and storage of these microbes. In this study, we present a simple protocol for optimizing cryopreservation of non-model organisms in laboratory settings using conventional chemicals. Our initial motivation for this observation was to discover a cryoprotection agent for independently cultured *Mycetohabitans* spp., a fungal endosymbiont. We tested several common bacterial cryoprotection agents like glycerol, bovine serum albumin (BSA), and dimethyl sulfoxide (DMSO) over an ultralow freeze-thaw cycle to determine an adequate method of cryoprotection for assorted bacteria. We observed different recovery rates across bacterial species and cryopreservation methods, and identified cryoprotectants that reliably resulted in viable bacteria for each of the strains tested. We present this as a resource for those working with other fastidious and host-associated bacteria that may be missing effective cryopreservation methods.

## Importance

The ability to cryopreserve bacteria is important for optimizing laboratory procedures, preserving strains that have been genetically manipulated, and growing fresh cultures of microorganisms without *in vitro* evolution from serial subculturing. There are several known cryoprotection agents of bacteria but there are limited accessible studies that collect these together and screen them for effectiveness with new bacteria studied in laboratory settings. With several fastidious and host-associated microorganisms emerging as new model systems, we aim to generate a resource for determining long term storage solutions for novel organisms of interest.

## Observation

Advancements in sequencing and metabolomic profiling have improved our ability to identify and culture previously intractable bacteria. In parallel, there has been an increased push to study and manipulate more environmental microbes, expanding beyond the typical model systems (Elmore et al. 2023; Jensen 2025; Salcher et al. 2025). Long-term storage, typically through cryopreservation, is vital to studying bacteria in the laboratory environment. Microbes that typically rely on a host, or those that occupy specified niches, may not be amenable to ultralow freezing except when exposed to specific cryoprotectants (Hasan et al. 2018; Guo and Weng 2020; Kihika et al. 2022). For example, bacteria belonging to the *Mycetohabitans* genus inhabit fungal mycelia and were initially described as “erratic” for culturing and storage (Partida-Martinez et al. 2007). We repeatedly observed that *Mycetohabitans* bacteria do not consistently survive the freezing process in the typical 20-30% glycerol storage that many related gram-negative bacteria recover from, reducing the tractability of the organism for genetic studies (Hasan et al. 2018). However, *Mycetohabitans* spp., like other specialized symbionts and host-associated bacteria, are important to study for their role as endosymbionts and the creation of mutant strains for phenotyping experiments requires long-term storage strategies. Thus, we set out to identify a reliable method of cryopreservation for *Mycetohabitans* spp. Recognizing that a simple protocol for cryoprotection screening in a multiwell plate format that employs chemicals typically found in a microbiology lab would be a useful resource for the research community, we expanded our work to show how our method can be used to identify effective cryoprotectants for cultured bacteria from multiple niches.

Cryopreservation is a common microbiological technique that works by halting cellular metabolic processes by cooling cells to a low temperature (Louis et al. 1994; Hubálek 2003). Cryoprotection agents prevent formation of ice crystals within cells to protect cell structure and functionality of cell components (Jiang et al. 2023). Without cryopreservation methods, ice crystal formation can rupture the bacterial cell membrane or lead to fatally high concentration of the remaining liquid phase water within cells (Calcott 1986). We compiled a list of known cryoprotectants that have been used for long-term storage of bacteria and are readily available within a typical microbiology laboratory (Table 1). The typically-used permeating agents such as glycerol and dimethylsulfoxide (DMSO) form hydrogen bonds with water thereby lowering its freezing point and protecting cell membrane from rupturing (Hubálek 2003). Skim milk and bovine serum albumin (BSA) may act as a buffer and protect cells from damage by freezing amorphously around it (Cody et al. 2008; Tapias et al. 2013). Sucrose has been found to also create a protective matrix around cells, and reduce ice crystal formation (Li et al. 2010). These cryoprotection agents were screened by themselves and in mixtures to determine whether a combination would increase recovery. Additionally, sodium glutamate (SG) was tested in combination as it is typically used in conjunction with another cryoprotectant to protect cell envelope integrity by providing preferential hydration (Chen et al. 2023; Di et al. 2023).

**Table 1.** Cryoprotection Agents Used for various Microorganisms.

| Table 1. Cryoprotection Agents Used for Various Microorganisms |  |  |
| --- | --- | --- |
| Cryoprotectant | Sample Bacteria | Citation |
| Glycerol | *Various microbes* | Kim et al., 2009, LaBauve and Largo, 2012, and Trisiswanti et al., 2023 |
| Skim Milk | *Various microbes* | Cody et al., 2008 |
| Bovine Serum Albumin | *Azotobacter* sp. | Rojas-Tapias et al., 2013 |
| Sodium Glutamate | *Lactobacillus delbrueckii* | Fonseca et al., 2003 |
| Dimethyl Sulfoxide | *Enterobacterales* | Tutrina and Zhurilov, 2024 |
| Sucrose | *E. coli* | Louis et al., 1994 |

For each bacterium of interest, turbid liquid cultures were grown from three separate individual colonies to act as biological replicates. A multiwell plate was set up to screen all cryoprotection agents at one time for each bacterium assessed. Each turbid bacterial culture was washed and resuspended in 10 mM MgCl2 to an optical density at 600 nm of 0.7 then 100 µL was dispensed in triplicate per cryoprotectant mixture tested to create three technical replicates per biological replicate; 100 µL of 10 mM MgCl_2_ was dispensed in control wells. Stock solutions of cryoprotection agents were added until all wells had a final volume of 200 µL with the final concentrations listed in **Figure 1**. Each 96-well plate was then placed directly in an ultralow freezer to be stored at -80°C for 30 days. After being frozen, each plate was thawed and 50 µL aliquots were transferred into a new 96-well plate with 150 µL of Lysogeny Broth or King’s B Broth. For our experiments, bacteria were grown in a LogPhase600 Plate Reader (Agilent Biotek) at 30°C for 24 hours for faster growing bacteria or 76 hours for *M. endofungorum* which has a slower growth rate, at which time the optical density at 600 nm was measured. Plate reader settings, times, and temperatures could be adjusted based on the organism of interest and availability of equipment. An ANOVA statistical test was applied to differentiate efficacy and we used a Tukey’s HSD test for comparisons of cryoprotectant mixtures for each bacterium.

**Figure 1.**
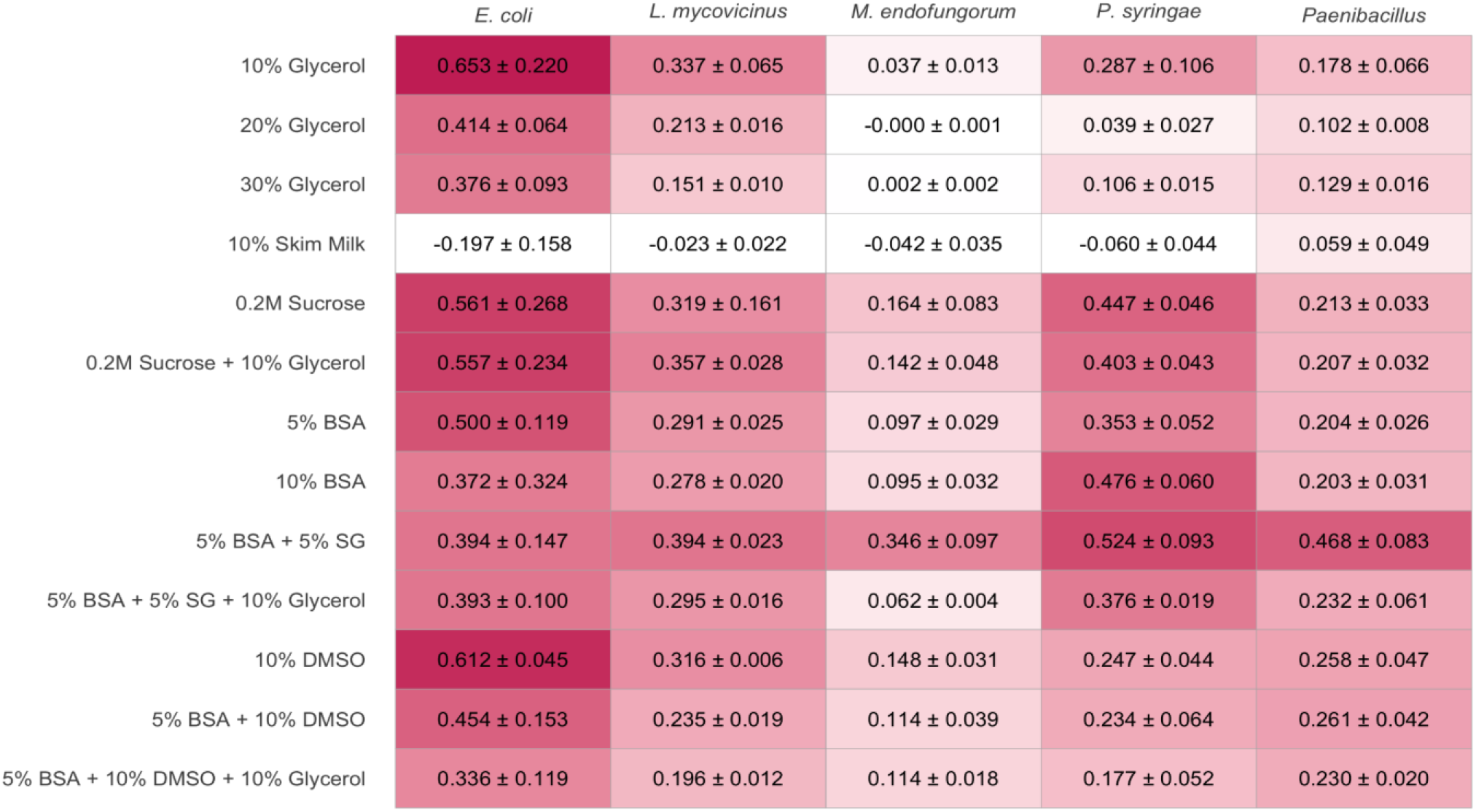
Bacterial recovery (mean+-sd) from cryopreservation at -80°C with the listed final concentrations of cryoprotection agents after one month at -80C. BSA, bovine serum albumin; SG, sodium glutamate; DMSO, dimethyl sulfoxide. Species tested were Escherichia coli S17-1, Luteibacter mycovicinus 9143, Mycetohabitans endofungorum B13, Pseudomonas syringae DC3000, and Paenibacillus sp. CB74. Growth was measured at 76 hours (Mycetohabitans) or 24 hours (all others) post inoculation.

As proof of concept, we tested the cryoprotectants on conventional laboratory *E. coli* (strain S17-1) and the model gram-negative plant pathogen *Pseudomonas syringae* strain DC3000. Our example host-associated bacteria for testing were the endohyphal bacterium *Luteibacter mycovicinus* strain 9143 and *Mycetohabitans endofungorum* strain B13. Our final inclusion was a novel *Paenibacillus* sp. CB74 that was recently isolated from fungal hyphae within our laboratory (unpublished) to represent more bacterial diversity.

Effective cryoprotection varied across all species in this sample set (**Figure 1**). Note that some resulted in negative average values when accounting for the blanks, but these were statistically indistinguishable from zero. In *E. coli*, we found that each cryoprotectant was adequate for cryopreservation with the exception of 10% skim milk (p < 0.001). *L. mycovicinus* also had a variable range of effective cryoprotection agents but showed minimal growth when preserved in 10% skim milk (p < 0.001). *P. syringae* was viable across a range of cryoprotectants but 20% glycerol, 30% glycerol, and 10% skim milk were least effective (p value < 0.001). In *M. endofungorum*, we saw that 5% BSA + 5% SG was the most effective cryoprotectant (p < 0.001). The inclusion of glycerol with BSA and SG reduced the viability of *M. endofungorum*. Finally, the novel *Paenibacillus* sp. showed robust recovery across cryoprotectants but 5% BSA + 5% SG was the most efficient (p<0.001).

As expected, we observed that recovery varied based on the cryoprotection agent used across different species. Some bacteria recovered from any tested cryoprotection mixture, but the growth post-freeze-thaw differed across conditions. In contrast, our primary bacterium of interest, *M. endofungorum*, was only recovered when one of the cryoprotection agents used was BSA and generally did not recover well from mixtures containing glycerol. We observed the most robust growth from the BSA and SG mixture for *M. endofungorum*. Despite being published as a viable alternative (Cody et al. 2008), skim milk was the least effective cryoprotectant tested. We found it was difficult to sterilize and maintain a homogenous liquid, with significant differences observed when we altered settings (sterilization time and volume), which suggests different outcomes could be achieved by modifying those. In our study, a combination of BSA and sodium glutamate worked effectively across organisms, though it was not necessarily the best option across all. The variation observed highlights the need for this simplified method for cryoprotectant screening, thus we present this useful guide for optimizing cryopreservation of novel microorganisms in laboratory settings.

## Acknowledgements

The authors would like to thank the #NewPI Slack community for their help in identifying cryoprotection agents to test and the Graduate and Postdoctoral Writing Center at UNC Charlotte for providing writing support on this manuscript.

